# Coastal Marine Mammal conservation using thermal imaging-based detection systems

**DOI:** 10.1101/2023.08.25.554754

**Authors:** Sebastian Richter, Harald Yurk, Alexander Winterl, Elly Chmelnitsky, Norma Serra, Patrick D. O’Hara, Daniel Zitterbart

## Abstract

Marine mammal habitats in coastal regions are increasingly threatened as anthropogenic use of these areas increases. Vessel traffic from commercial and pleasure craft leads to increased underwater noise and risk of marine mammal vessel-strike. Mitigation measures advised by competent authorities, such as static and dynamic speed restriction zones or area closures, must be based on reliable data to be justified and find widespread adoption. Thermal detection systems can augment currently employed monitoring approaches such as passive acoustic monitoring or marine mammal observers. Thermal imaging whale detection systems are cost-effective, can be operated around the clock and do not rely on the animal to vocalize, hence have a different availability bias than passive acoustic monitoring. It has been shown in previous studies that thermal detections based on a pronounced blow is possible for larger baleen whales. In this study, we evaluate how effective thermal imaging base whale detection systems are for detecting smaller marine mammals, with a focus on killer whales in Salish Sea.

## Introduction

In animal conservation management one of the important questions is how much time animals spend within a suitable habitat and how they move within that habitat and between suitable habitats. The problem becomes exacerbated by the behavior of highly mobile species that may use a variety of habitats for different purposes over the course of a season or year. Acoustic recorders and cameras have been used in both terrestrial and marine environments to detect and track animal movements over shorter and longer time scales (Yurk *et al*., 2010; Kalan *et al*., 2016; Evans, Mosby and Mortelliti, 2019; Palmer *et al*., 2019; Lahoz-Monfort and Magrath, 2021). Acoustic recorder-based systems have certain advantages over camera-based systems, including that they are less affected by weather conditions. They are, however, affected by ambient noise which reduces such systems’ detection range. Furthermore, passive acoustic monitoring systems suffer from the availability bias that animal must vocalize to be available for detection. Underwater soundscapes are dynamic and sound propagation is highly variable and is influenced by environmental factors, such as bathymetry, water temperature, and pressure which affect the underwater sound speed. Detection ranges may vary among locations and between different seasons. Additional anthropogenic noise (e.g., boat noise) may mask the signals and thereby further reduce detection range. As a result, whale monitoring often relies on a combination of visual observation and acoustic monitoring. Visual monitoring using cameras or observer sightings may often be limited to specific seasons when light and weather conditions are good and require dedicated and experienced observers, while passive acoustic monitoring requires trained acousticians, appropriate analysis software, and specialized recording equipment. Both methods can become very costly when there is a need to detect and track whales in near real-time. Near real-time detection and tracking, however, is sometimes necessary to mitigate threats that result in injury or death of individuals, such as vessel strikes or exposure to high sound energy such as sound created during seismic surveys, pile driving and sonar operations (Verfuss *et al*., 2016, 2018).

In recent years thermal-imaging based whale detection systems have been added to the whale detection toolbox. Such systems can be operated at day and night, are not affected by sound masking, do not require the marine mammal to vocalize, and can operate without a human observer. Artificial intelligence driven whale detection using thermal imagery (TI) is now semi-routinely used to detect cetaceans (Zitterbart *et al*., 2013). Albeit range limited, the method has been shown to detect large whales at distances similar to human observers (Zitterbart *et al*., 2020). An assessment of the performance of such detection systems has been conducted for large whales (Smith *et al*., 2020; Zitterbart *et al*., 2020). A study performed in Haro Strait (Graber *et al*., 2011) showed that killer whales can be detected based on the thermal signature of their body and their blow, however, the long-term performance of such systems has not been assessed for small to mid-sized odontocetes. In this study, we evaluated the covariates that influence detectability and the feasibility to use TI systems to automatically detect killer whales in the inshore waters off the coast of British Columbia, Canada, called the Salish Sea. Covariates considered in this study are the angular resolution of the thermal imaging camera systems, elevation of the observation location, species, and marine mammal group size.

### Killer Whales (Orcinus orca) of the Northeast Pacific including the Salish Sea

Three ecologically distinct killer whale forms or ecotypes inhabit the Northeast Pacific, *Offshore, Bigg’s (formerly transient) and resident killer whales* (Parsons *et al*., 2013; Ford and Nichol, 2014). The Offshore Killer Whale tends to occur in pelagic waters off the coast and forages on fishes including sharks. Within the Salish Sea typically only Bigg’s (transient) and resident killer whales occur. Bigg’s Killer Whales (Bigg’s whales) occur both in nearshore habitats including channels, inlets, and bays and in pelagic waters offshore specializing on foraging on marine mammal prey including pinnipeds, dolphins, and porpoises. Bigg’s whales encountered in the Salish Sea often travel in groups of three to five animals when foraging, but sometimes also form larger social groups (Baird and Dill, 1996; Baird and Whitehead, 2000). Resident killer whales (RKW) also venture into waters over the continental shelf but can be found fairly regularly in inland waters including channels, fjords, and bays which are common in the Salish Sea (Ford *et al*., 2017; Shields, 2023). RKW live in stable family groups consisting of a female and all her female and male offspring and these groups can comprise up to four generations. Individuals tend to leave these groups which are called *matrilines* only temporarily and for very short periods (hours). Closely related matrilines (pods) spend more time associating than distantly related matrilines. This typically leads to large groups of animals (> 15 animals) traveling together. When resident killer whales are foraging on salmonids with a clear preference for the largest salmon in the Pacific, the Chinook salmon (Ford and Ellis, 2006; Hanson *et al*., 2010), they often spread out over as large an area as staying in contact acoustically allows. There are two distinct resident killer whale populations/communities along the coastlines of British Columbia and adjacent states of the USA called Northern and Southern Resident Killer Whales (NRKW and SRKW). SRKW are the more commonly encountered RKW in the Salish Sea along with Bigg’s whales. The differences in geographical distributions, social structure, and foraging behavior led to differences in occurrence, encountered group size and movement behavior that influence detectability of each ecotype.

#### Conservation concerns

Both RKW populations are considered at risk of extinction due to their low numbers and the SRKW, which only contain 73 members as of July 2022, are designated as an endangered population in Canada and the USA (NOAA, 2005; DFO, 2018). The SRKW face a number of threats to their survival; declining chinook salmon stocks and availability as prey, acoustic and physical disturbances primarily from vessels including the risk of fatal vessel strikes, oil spills and other contaminants (Cosewic, 2008).

SRKW comprise three pods, called J, K and L pod (Fisheries and Oceans Canada, 2011). While the three pods disperse at times especially during fall and winter, they stay mostly near the coast and travel between Oregon, Washington and British Columbia (Ford *et al*., 2017; Rice *et al*., 2017; Emmons, Hanson and Lammers, 2021).

The Canadian recovery plan for SRKW (Fisheries and Oceans Canada, 2011) lists a number of actions including those that address physical disturbance, which had been identified as a threat to the recovery of the population (Cosewic, 2008). Fisheries and Oceans Canada also specifically recognizes the threat posed by injurious and fatal ship strikes to the survival of the SRKW population, *“The very small size of the SRKW population and the low numbers of prime age males and females that support the reproductive potential and genetic diversity of the population means that a threat that could remove one animal will have significant consequences.”* (Fisheries and Oceans Canada, 2011).

### Reducing the risk of ship strikes through near real-time detection

The potential of ship strikes having negative population consequences for whales including SRKW prompted the Department of Fisheries and Oceans Canada (DFO) to create the Whale Detection Collision Avoidance (WDCA) initiative, that aims at reducing the risk of physical disturbances including vessel strikes through the development and testing of near real-time (NRT) whale detection and tracking technologies. The outcome of the work of the initiative was to provide DFO with methodologies that allow NRT detection and tracking of whales to initiate appropriate management actions such as issuing mariner alerts of whale presence. The initiative was funded by Canada’s Ocean Protection Plan^1^. In Canada’s Pacific region the initiative focused on methods to detect SRKW but the methodologies were evaluated on their effectiveness to detect all marine mammals in NRT. To reduce the risk of ship strikes NRT detection and tracking of whales including tracking of the three SRKW pods would be ideal but is difficult to achieve due to the high mobility of the animals and the fact that not all pods occur in the same place at the same time. One way to address this problem is to monitor whale movements in locations with high overlap of whale and vessel occurrence and test the effectiveness of detection and tracking technologies in those locations with greater likelihood of potential ship strikes.

In this study we present the results of testing thermal-imaging based automated systems to detect killer whales in narrow waterways within the Salish Sea, Active Pass and Boundary pass, with a potentially high risk of ship strikes (see Error! Reference source not found.). Active Pass and Boundary Pass are major thoroughfares for large commercial vessels, ferries, and smaller non-commercial vessels. SRKW and humpback whales frequently use these topographically constrained waterways to travel between southwestern and northeastern sections of the Salish Sea.

## Material & Methods

### Location

This study includes data collected at two field sites located in the Salish Sea, British Columbia, Canada. One on Galiano Island monitoring Active Pass, a narrow waterway that is used regularly by ferries and other commercial and recreational vessels. A second one close to East Point on Saturna Island overlooking a section of Boundary Pass that includes the international shipping lanes connecting the Port of Vancouver with the Pacific Ocean (see Error! Reference source not found., Table **1**). Active Pass is considerably narrower at the camera site (1500m) than Boundary Pass (5400m).

**Table 1.** Field sites where the thermal whale detections systems were deployed.

| Location | Latitude | Longitude | Elevation (m, MSL) | Data collection period | Note |
| --- | --- | --- | --- | --- | --- |
| Galiano Island, Sturdies Bay, Active Pass | 48° 52' 35.609" N | 123° 18' 53.507" W | 14 | 7 June 2019<br>31 March 2021,<br>22 months | 3 cameras,<br>6.4° bearing 93°<br>12.4° bearing 114°<br>25° bearing 131°<br>max visible range<br>1300m |
| Saturna Island, East Point, Boundary Pass | 48° 46' 51.658" N | 123° 3' 6.530" W | 15 | 4 August 2020<br>31 March 2021,<br>8 months | 1 camera,<br>12.4° bearing 172°<br>max visible range<br>5500m |

The Galiano Island system was installed on top of the Sturdies Bay ferry terminal operated by BC Ferries. Multiple thermal imaging and visual cameras were mounted at an elevation of 14m (above Mean Sea Level, MSL) and pointed eastward across Active Pass. The system was operated with a single thermal imaging camera (6.2° field of view) from June 2019 to November 2019 and two thermal imaging cameras (12.4° and 25° field of view respectively) from November 2019 to March 2021 for a total of 22 months.

The Saturna Island system was installed on a property belonging to a member of the Saturna Island Marine Research and Education Society (SIMRES). A single thermal imaging camera (12.4° field of view) was mounted at an elevation of 15m (above MSL) pointing southwards across a section of Boundary Pass towards Skipjack and Waldron Islands.

### Thermal (IR) Imager

Thermal imaging data was acquired using an uncooled V0x microbolometer (A65, FLIR Systems, USA) in a spectral range of 7.5 – 13 µm (long wavelength infrared, LWIR) with three different lenses with horizontal field of views of 6.2°, 12.4° and 25° and a resolution of 0.01°, 0.02° and 0.04° per pixel (focal lengths of 100mm, 50mm, 25mm). Cameras were deployed on elevated positions between 14m and 15m in environmentally protected housings (Orca, autoVimation, Germany). Thermal data was acquired at 15 fps and evaluated in real-time (Zitterbart *et al*., 2013, 2020).

### Video (VIS) Camera for verification

At Sturdies Bay, the two thermal imagers were paired with two 5.1Mpx RGB cameras (GT2460 Color, Allied Vision, Germany) with a 25mm lens resulting in a 20° horizontal field of view (LM25JC5MM-IR, Kowa, Japan) to verify IR detections visually. The lens aperture was dynamically controlled via a P-Iris interface and focus was manually set to a fixed value. Color images were acquired at 7 fps and stored for each automatic detection during daylight hours to allow for species verification of thermal imaging detections.

### Algorithm and performance metrics

The detection and classification algorithm used to evaluate the thermal data stream in real time is based on (Zitterbart *et al*., 2013). Thermal anomalies caused by either blows or surfacing events, a whale exposing the whole body (breach) or parts of the body (back, dorsal fin or fluke), are automatically detected using their local heat contrast to the surrounding water surface and processed by a machine learning based classification system to determine the probability of a whale detection. Likely whale detections are then uploaded for validation by an operator.

We evaluated the automatic detection performance using a distance dependent detection function. A detection is defined as one individual sighting of a blow or surfacing event, implying that one individual crossing the field of view can be detected multiple times.

Automated detections were provided to a human operator for validation as short 6 second video snippets (thermal and visual if available). In addition, the species was annotated if an identification was possible based on the blow and body signatures in thermal and visual recordings. Data examples for a humpback whale, killer whale, pinniped and porpoise detection are shown in Figure **1**.

**Figure 1.**
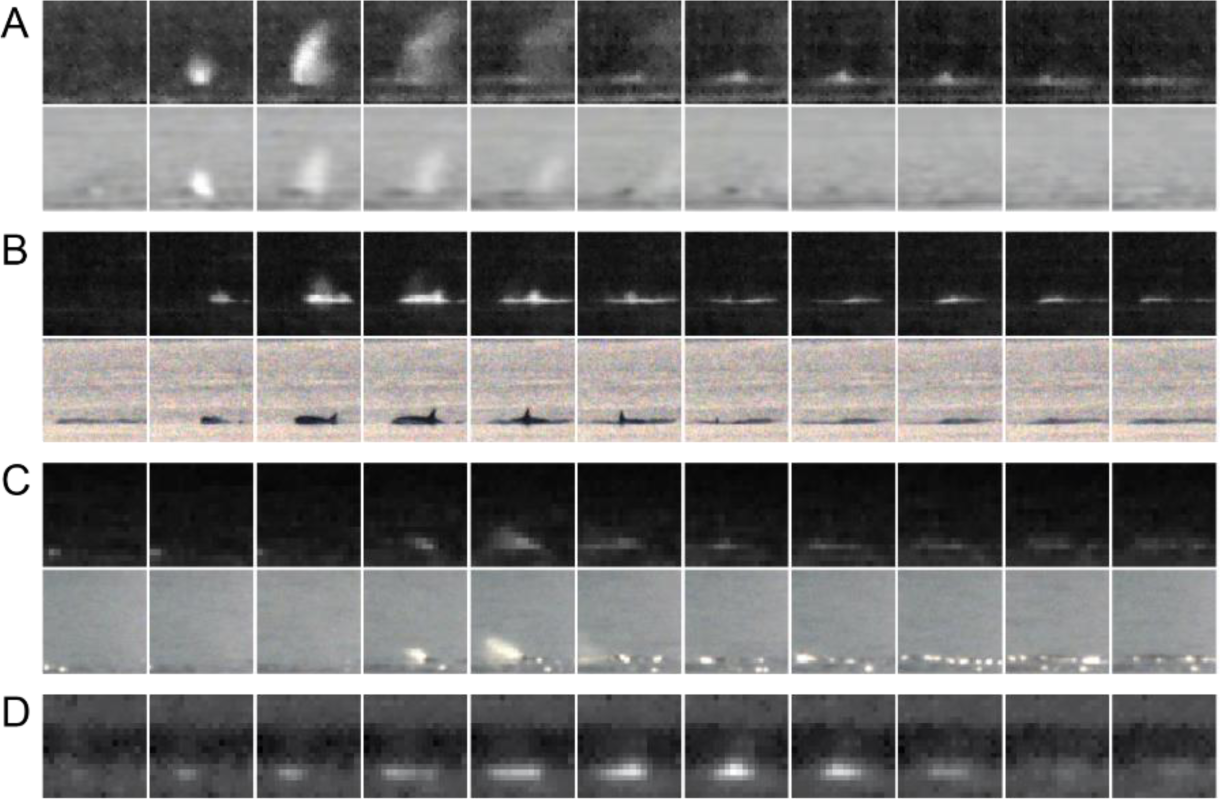
Thermal and complementing RGB data (if available) of automated detections by the systems in Sturdies Bay for multiple species, humpback whale (a), killer whale (b), seal (c) and harbor porpoise (d). Time delta between frames is 0.4s, with the exception of the porpoise detection (d) which has a time delta of 0.2s.

Killer whales travel as groups that contain males, females, and juveniles. The focus of the study is to detect at least one of the animals in a group but preferably more than one. Based on the validated data we generated distance dependent detection functions, similar to the data processing for a point-transect distance sampling protocol (Buckland *et al*., 2004). Under the assumption of equally distributed whales in the observation area, the number of detections grows linearly with distance (Buckland *et al*., 2001), as the area of a ring or distance bin increases linearly with increasing distance. At some point the detection functions will begin to decrease as the system begins to miss detections. The location of the detection functions peak characterizes the *reliable detection distance*, the range at which all whale cues are detected. The detection with the longest distance describes the *maximum detection distance*, the maximum range at which the system was able to detect a whale.

## Results

Assuming a uniform distribution of marine mammals, according to point transect distance sampling approach (Thomas *et al*., 2010) we would expect a linearly increasing number of detections with increasing distance due to an increase in monitored area. However, with increasing range the signal-to-noise ratio of the thermal whale signature is decreased, and the detection system will miss whales’ surfacing, thus probability of detection will decrease. The location of the peak of the detection function characterizes the reliable detection range (Zitterbart *et al*., 2020; Baille, LMR and Zitterbart, DP, 2022) which depends on several parameters such as the size of the blow and visible body area of the target species, the camera resolution and elevation of the camera. Detection functions for killer whales at the two field sites and lens configurations are shown in Figure **2**.

**Figure 2.**
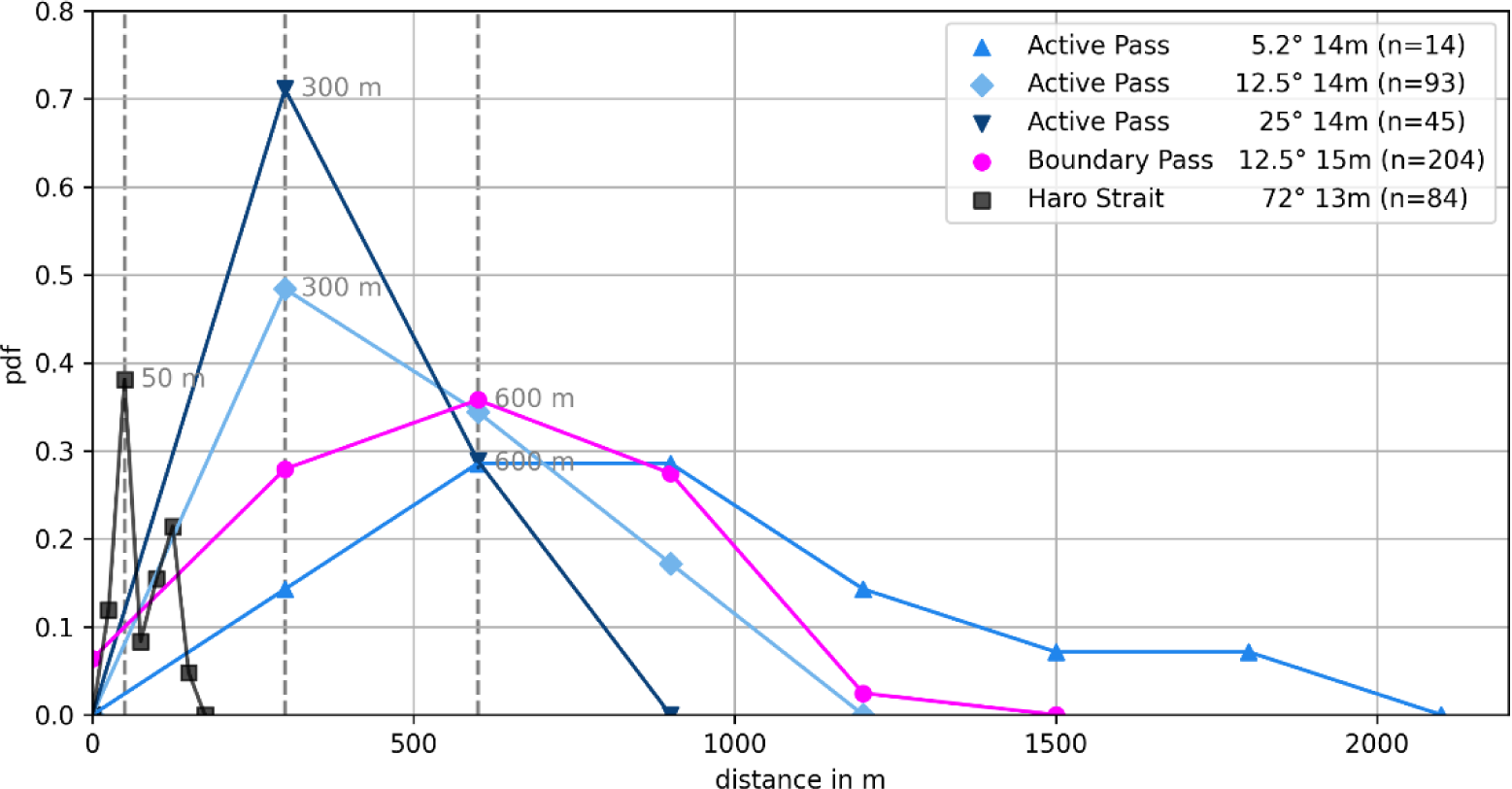
Detection function for killer whales on different locations (Sturdies Bay at 14m height, and Saturna Island at 15m height) and angular resolutions (horizontal field of views 6.2°, 12.4°, 25°, 72°) in comparison to a previous study in Haro Strait (Graber et al., 2011).

At Sturdies Bay, reliable detection distance was approximately 300m for both 12.5° and 25° FOV cameras. However, only the 12.5° lens had some detections in the 900m range. Due to the orientation of the camera across Active Pass, detections further than 1300m were not possible due to the limited width of Active Pass. The 5° camera was pointed further north (bearing 93°), towards the northern end of Active Pass and into the Strait of Georgia, the maximum number of detections was at 600m and furthest detections ranged up to 2100m. The Saturna Island camera (12.5°) pointed south across Boundary Pass and showed reliable detection distances of 600m and the furthest whale detections exceeded 1300m.

We further analyzed species specific detection functions as shown in Figure **3**. In Active Pass humpback whales (*Megaptera novaeangliae)* have been detected at distances between 400m to 2150m (RDD: 500m). Detections beyond 1300m originated from the first deployment phase using the 5° lens pointing towards 93° east. Killer whales have been detected at distances from 300m to 1900m (RRD: 500, Figure **3**). Northern Minke whales *(Balaeonptera acutorostrata)* have been detected at distances from 450m to 550m. Pinnipeds as well as Harbor porpoise *(Phocoena phocoena)* detections only occurred in close proximity to the camera between 100m to 700m (RDD: 200).

**Figure 3.**
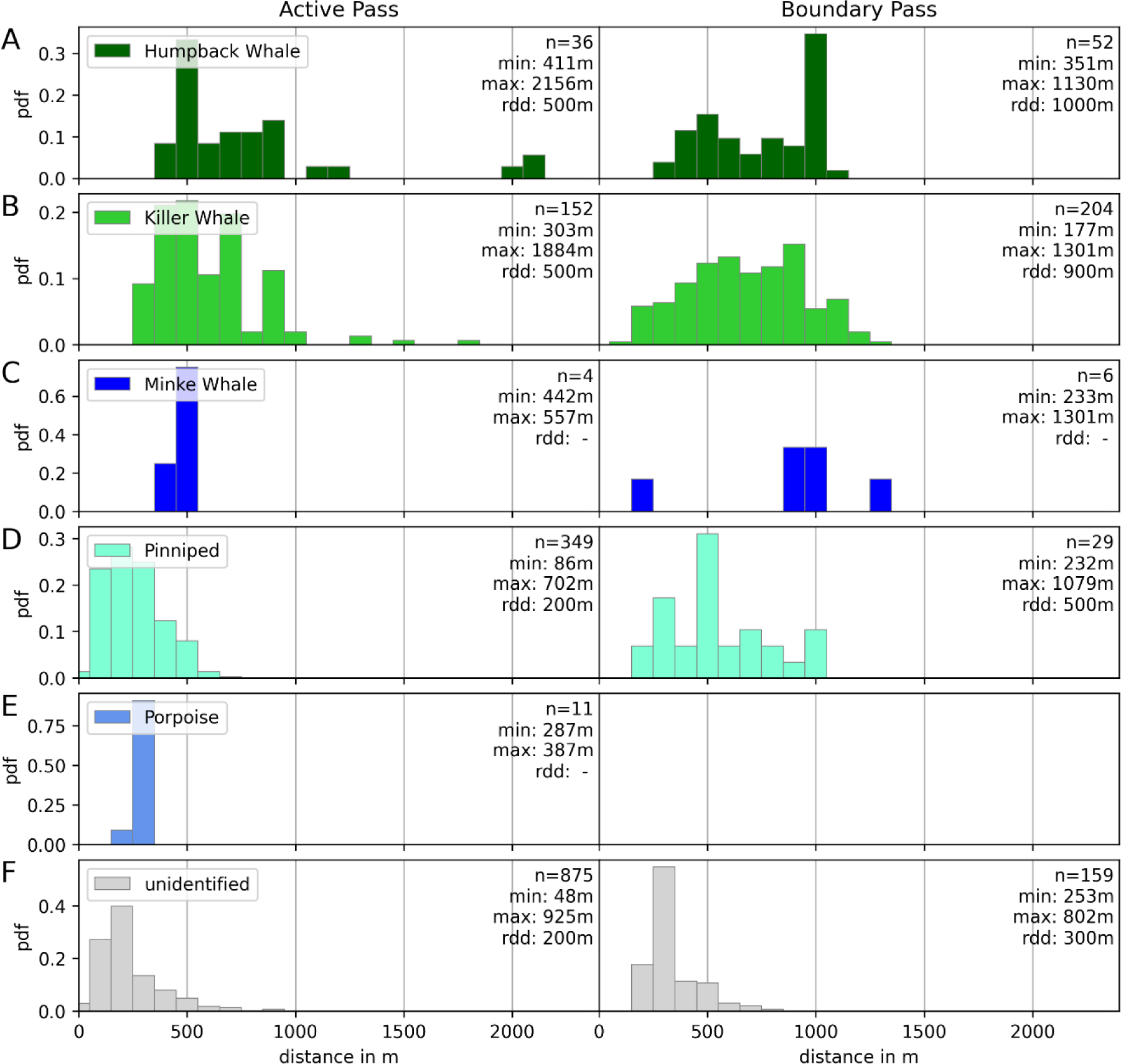
Detection functions by location Active Pass (left) and Boundary Pass (right) and species (humpback (A), killer (B), minke whale (C), pinniped (D), harbor porpoise (E) and unidentified small animal (F)). Detections are pooled for all three lens models for Active Pass.

At Boundary Pass, humpback whales were detected at distances between 350m to 1100m (RDD: 1000m) and killer whales between 170m to 1300m (RDD: 900m). We detected three events that are most likely minke whales on two different days at 170m and 1300m. Pinnipeds were detected between 200m and 1300m and there have been no detections which could be identified as harbor porpoise.

At both locations, most detections that could not be assigned to a species were close to the camera and most likely belonged to small marine mammals, fish, or larger diving birds.

Continuous observation throughout day and night allowed assessing species occurrence throughout the whole observation period. The distribution of automated detections over time are shown in Figure 4. Generally, detections are rare (on average 0.62 detections per day across locations for humpback, minke, and killer whales), with long periods without any detections (mean: 11 days, median: 4 days, maximum:124 days during summer 2020).

**Figure 4.**
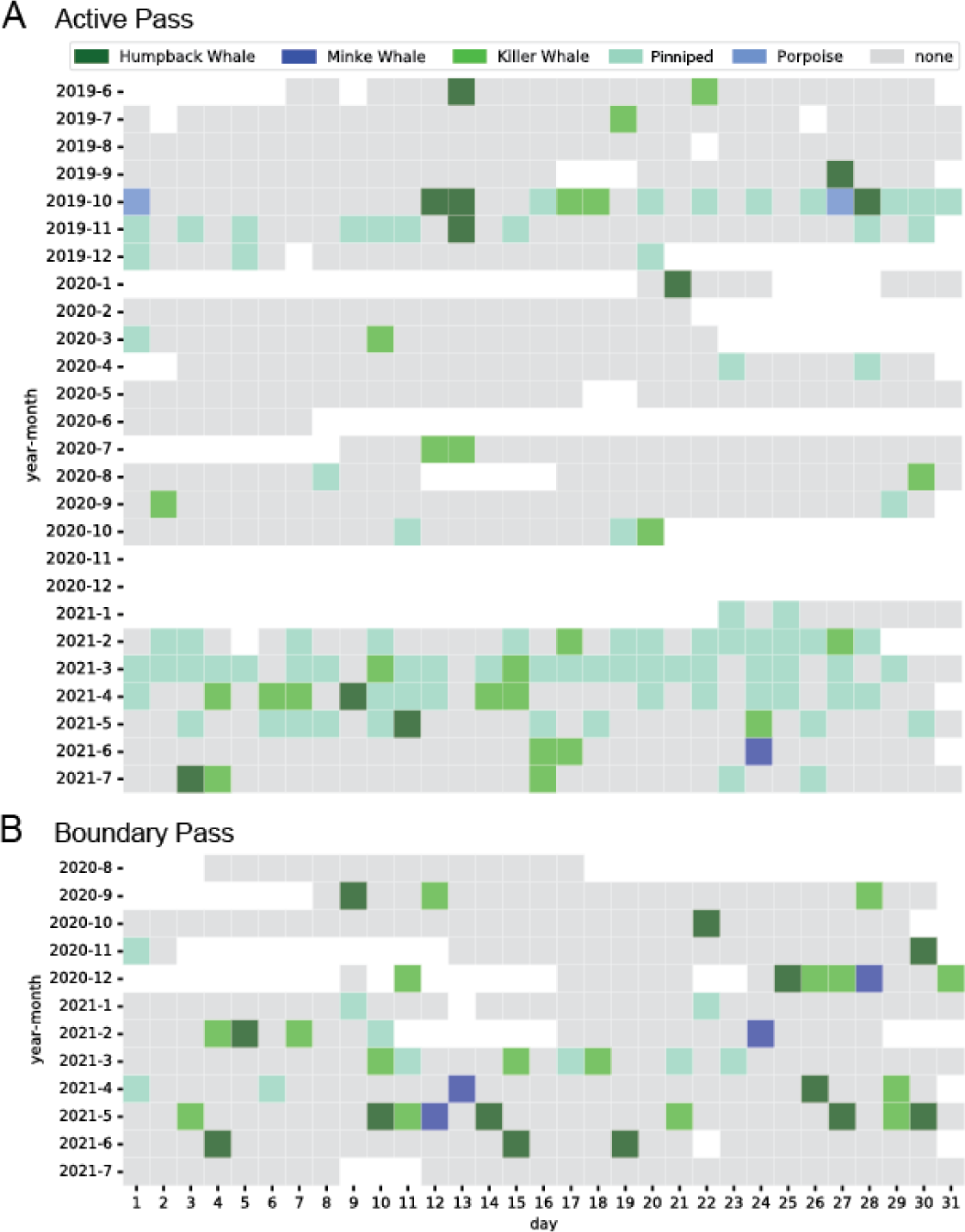
Detections by species throughout the deployment period. Days without data due to power loss, lost connectivity, or lack of access to the system due to COVID-19 travel restrictions, are shown in white. In case of multiple species detections in one day the order of ranked occurrence is humpback whale, minke whale, killer whale, pinniped and porpoise.

For a better description of species occurrence, we define an encounter as multiple consecutive detections of the same species within a period of less than 10 minutes. In Active Pass (Figure 4 A) we automatically collected 36 humpback whale detections in 13 encounters, 152 killer whale detections in 26 encounters, 349 seal detections in 155 encounters and 11 harbor porpoise detections in four encounters. Humpback whale detections have been most frequent during the first operation period from June 2019 to November 2019 where one 5° field-of-view thermal camera was used to monitor the northern entrance of Active Pass. A smaller camera angle provides a greater detection range, and the whales may not have been close enough to Sturdies Bay to be detected by the camera with the wider angle. Killer whale detections were more evenly distributed across the observation period. Seals were frequently identified during the winter months and porpoises have only been identified on two days during the observation period. Interruptions in the operation of the systems have been mainly caused by frequent multi-hour power outages or a lack of data storage space due to COVID-related travel restrictions, when data hard drives could not be replaced in time.

In Boundary Pass (Figure 4 B) the system made 52 humpback whale detections in 14 encounters, 204 Killer whale detections in 16 encounters, six Minke whale detections in four encounters and 29 seal detections in 14 encounters. Interruptions in the operation have been mainly caused by power outages.

## Discussion

### TI detection for KW

This study was specifically designed to evaluate the performance of TI detection systems for the highly endangered SRKW population. In contrast to humpback whales, which were the target species in previous studies (Zitterbart *et al*., 2013, 2020; Smith *et al*., 2020)the blow of a killer whale is less pronounced, which makes it more difficult to detect. However, killer whales expose a large portion of their body including their characteristic dorsal fins which can be used to identify the species in thermal images (Graber *et al*., 2011). Hence, most killer whale detections are based on the body signature instead of the blow signature. A distinction between the two killer whale ecotypes (resident, i.e. SRKW or Bigg’s whales), however, is not possible based on the thermal imaging detection. Because of differences in the social and feeding ecology of the different ecotypes, more detections per encounter, could be indicative of the presence of SRKW due to the larger group sizes of SRKW (15-45 animals) versus Bigg’s whales (1-15 whales). However, group size alone is not a definitive distinction between RKW and Bigg’s whales.

A reduction of detection range due to adverse weather conditions such as fog, wind, and heavy rain as well as strong sunlight producing water surface glare has been previously discussed for large baleen whales across different ocean regimes, ranging from polar to tropical environments (Smith *et al*., 2020; Zitterbart *et al*., 2020) and similar negative impacts on killer whale detectability were expected.

### Location specific performance

The detection functions between the two locations are similar, as we would expect based on the similar camera elevation and lens models (see Figure **2**). The remaining differences are largely based on the observable ocean surface. While the Active Pass detection functions suggest a reliable detection range of only 300 meters, detections acquired by the same camera and lens model at Boundary Pass indicate a reliable detection range of 600 meters. This lower detection range in Active Pass is likely due the reduced observable surface area, which is more limited by the width of the pass at this site (1500m) than at Boundary Pass (5400m). The maximum detection range is further decreased by reflections of the opposite coastline on the water such as trees, that appear as bright objects in the thermal image and hence reduces the signal to noise ratio of the whale blow.

Furthermore, the similarity in detection functions for 12.5° and 25° field-of-view cameras in Active Pass can most likely be explained by preferred travel patterns. While the narrow field-of-view camera would be able to detect killer whales further away, e.g., close to the opposite shoreline, this does not result in a difference in detection functions due to a lack of individuals in this area as the assumption of equal distribution does not hold.

At Boundary Pass, we do not observe any detections farther than 1300m from the camera, which is less than half the width of the pass at the camera site. Killer whales may use bathymetric features to inform their travel paths and SRKW may travel less often in deeper waters which also are closer to the shipping lane in this area. Visual observations by human observers (L. Quayle, pers. comm.) indicate that SRKW often travel close to East Point when either entering or leaving Boundary Pass from or to the Strait of Georgia. Because the camera is located close to East Point and SRKW travel regularly within 2 kilometers of the shoreline when passing the camera they are likely to be detected by the camera. Bigg’s whales also tend to forage or search for their preferred prey (harbor seals) within 3 km of the shoreline and are less likely encountered at greater distances from shore (Ford, Ellis and Stredulinsky, 2013) but could occur on either side of the pass.

The temporal distribution of killer whale detections in Active and Boundary Passes indicates that the species can be present any time of the year. While this also holds true for humpback whales, the detection quantity appears to indicate a more seasonal use of the monitored areas. Days with detections in the middle of winter are fewer than they are for days in the spring, summer and fall. There is, however, a slight bias in the detectability of the two species, which leads towards a higher probability of detecting killer whales who travel in groups versus humpback whales who may travel as individuals. A whale has to surface in the FOV of the camera to be detected. A lower density of humpback whales in the winter may result in a higher rate of missed detections for this species.

#### Species other than killer whales detected by TI

While this manuscript focuses on killer whale detectability, the camera systems detected multiple other species (Figure 4) which we briefly discuss for completeness. Pinniped detections appear to show a seasonal distribution in Active Pass with higher occurrence during the late fall and winter. This may be due to the presence of Californian sea lions that seem to haul out at the southern end of Active pass during that time period. California sea lions migrate north from California in the fall and back south during the spring. Pinniped detections in Boundary Pass are also more frequent during late fall and winter. The data seem to indicate that sea lions are more likely detected by TI systems than the smaller Harbor seals which occur year round in both areas (Ford and Nichol, 2014).

Porpoise detections are overall sporadic in both areas which may be due to the small size of harbor porpoises (1.4 – 1.9 m), the predominantly occurring species in both areas (Ford 1984). Harbor porpoise detection range is limited to a distance of several hundred meters from the system.

Minke whales are also only detected sporadically by the TI system, which is likely due to the behavior of the species. Minke whales tend to travel and forage as single individuals in these waters (Ford and Nichol, 2014).

### Capability to detect rare events

The capability to detect a passing individual or group is defined by the chance to detect an individual at a specific distance, given by the detection function, and the number of detectable surfacing events. Figure **5** shows the fraction of encounters (y-axis) that consist of at least n detections (x-axis). Eighty five percent of killer whale encounters have more than one detected event. For humpback whales this is the case in 52% of all encounters.

**Figure 5.**
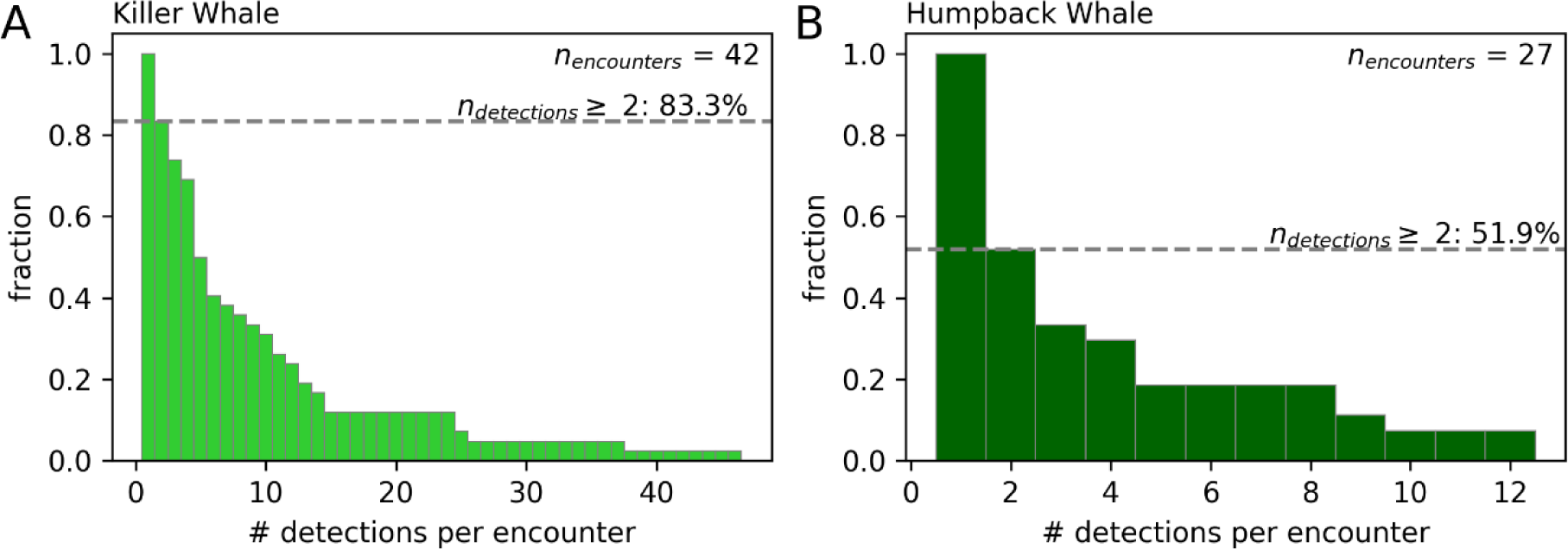
Number of detections per encounter (dt < 10 minutes) shown as complementary cumulative distribution function (CCDF) for killer whales (A) and humpback whales (B). Killer whale encounters consist of more than one detection in 83,3% of encounters (median 5 detections), humpback whale encounters consist of more than one detection in 51.9% of encounters (median 2 detections).

We can describe the chance to miss an encounter (*P*_*miss*_) given the distance of the event (*d*_*e*_) and number of events (*N*_*e*_) as one minus the probability to detect a surfacing event in a given distance to the power of the number of detected events per encounter:

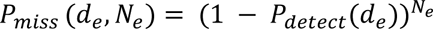

Even without a 100% chance of detecting an individual surfacing event at a given distance, the chance of missing an encounter with multiple surfacing events is small. The relation of detection probability for individuals and encounters with varying group size is shown in Figure **6**.

**Figure 6.**
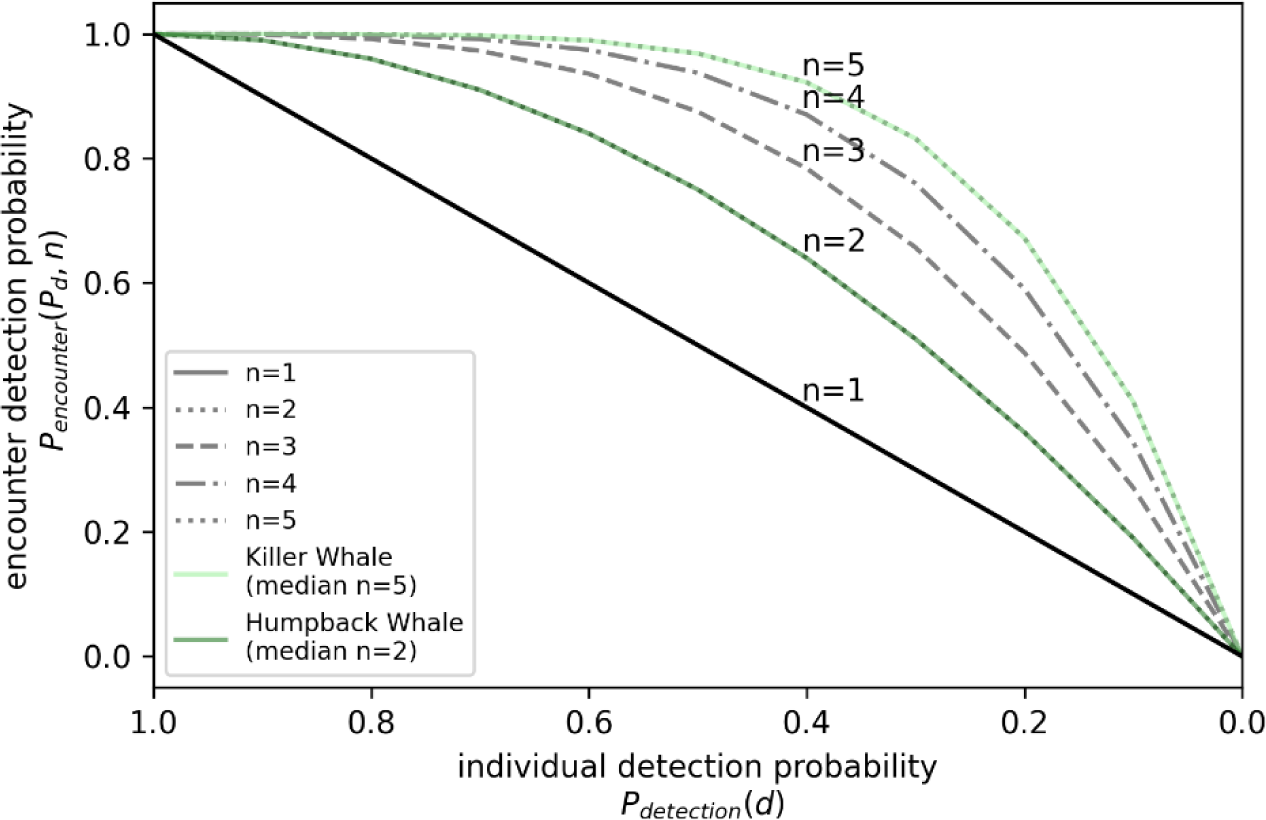
Comparison of encounter detection probability (1-P_miss_) over individual detection probability. The larger the encounter size the lower the probability of missing the encounter even at diminishing individual detection probabilities with increasing distance. The median encounter size for killer whales in the recorded data is five detections. Assuming an individual detection probability of 0.4 the probability to detect the encounter remains highly likely at 0.92.

In the context of collision avoidance and mitigation measures, especially when the target population is SRKW, missing an individual detection is not critical as long as the encounter is detected. Most SRKW encounters consist of more than one detection, hence missing an encounter even at distances exceeding the RDD becomes less likely with increasing number of detections per encounter. The risk of missing an encounter increases for species with small group sizes and fast traveling speed, as the chance for a blow or surfacing event to occur inside the field-of-view of the camera decreases. This can be mitigated by adding more cameras to increase the field of view where necessary.

### Effects of lens angle on coverage and spatial resolution

The choice of lens and the resulting field-of-view constitutes a trade off between coverage area and spatial resolution. While a shorter focal length offers a larger field of view and allows for the monitoring of a larger area (see Figure **7**A, 50mm lens with 12.5° horizontal field of view (left) and 25mm lens with 25° horizontal field of view (right)) the spatial resolution decreases proportionally (0.02° and 0.04° per pixel).

**Figure 7.**
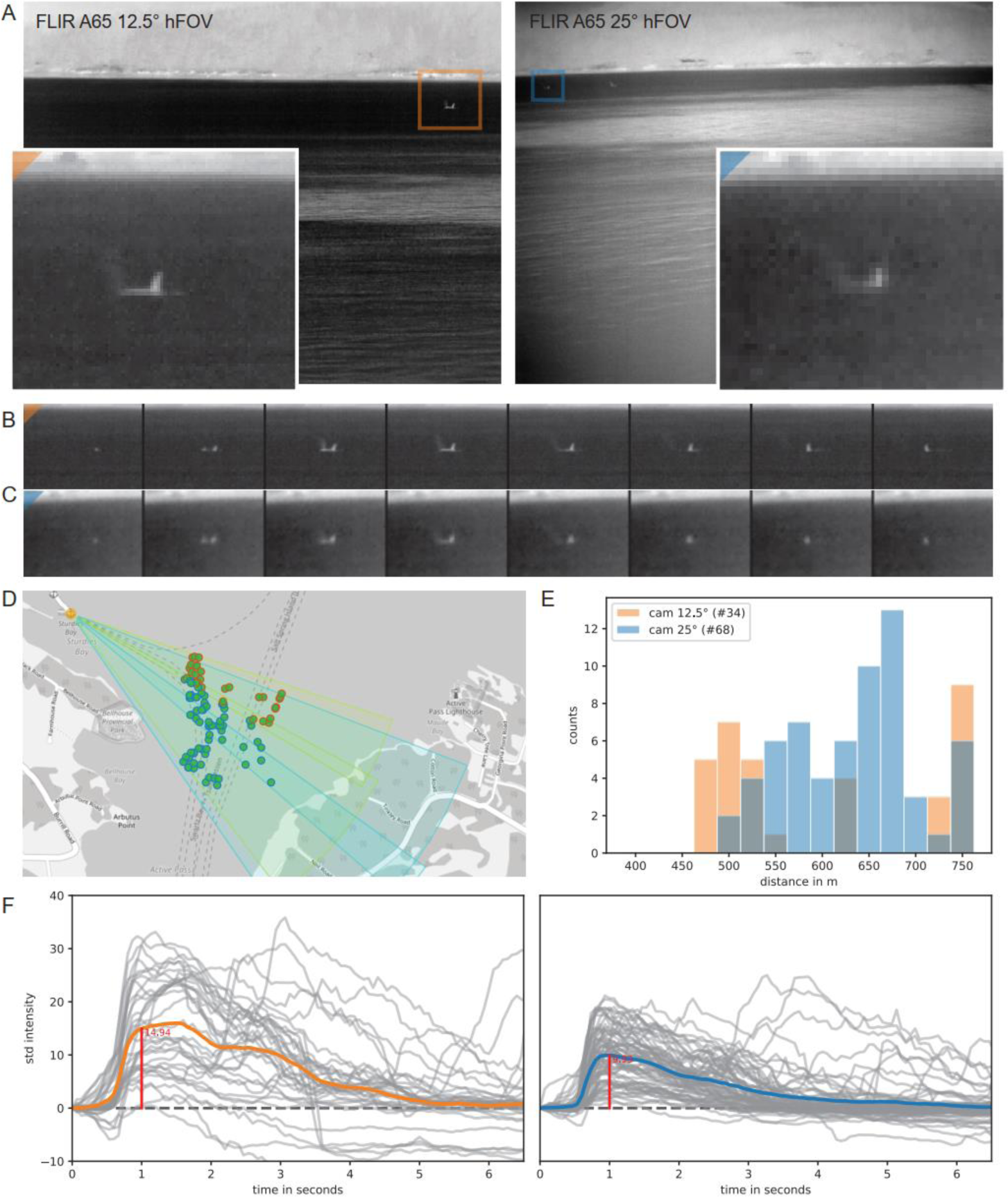
Compare FOV / Resolution (12° vs 25°) based on manual annotation of a Killer Whale encounter. The same surfacing/blow of a male Killer Whale at 550m recorded with both a 12.5° lens (A, left, marked in orange) and a 25° lens (A, right, marked in blue). The inserts illustrate the difference in spatial resolution. Time evolution of the thermal signal for both cameras with dt = 0.4s (B, C). Georeferenced location (D) of detections and distance of detection to the camera (E): 12.5° lens indicated in orange, 25° lens in blue. Estimate of the signal-to-noise ratio based on the standard deviation of intensity values over time in the area of a detection (F). Time evolution of each sample (gray) and mean value over all samples in the respective color. The average signal at t=1 is ⅓ lower for the 25° lens (9.93, blue) compared the 12.5° lens (14.94, orange)

The overlapping orientation of the field of view of our two cameras in Active Pass (Figure **7**D) enabled us to record the same surfacing event with two cameras with a 50mm and 25mm lens respectively. The loss in spatial resolution can be seen in the zoomed inset (Figure **7**A) and the time evolution of the surfacing event (Figure **7** B and C).

In addition to the loss in spatial resolution the signal to noise ratio deteriorates with increasing field of view. The smaller the spatial resolution, the more area is mapped to one pixel, and the more we average over and blur fine structures. As a result, the maximum thermal radiation measured decreases, for example when a pixel partially covers the whale body and partially the surrounding ocean, the resulting measurement will reflect a mixture of the thermal radiation emitted and reflected by the whale body and the ocean.

We can see this influence in our experiments. The annotated killer whale surfacing events are in a similar range to the camera (Figure **7**E), however we can see a clear reduction in measured thermal radiation relative to the background (Figure **7**F). The average received signal at t=1 (i.e., 1 second into the surfacing event, standardized for maximum signal) for the wider 25° lens (25mm) is 33% lower than the average signal for the narrower 12.5° lens.

In summary, a 50% wider field of view not only reduces the spatial resolution by 50%, in addition it reduces the signal strength by 33%. Selecting a fitting camera and lens model for the desired detection range and location is critical.

### Comparison to other approaches

Thermal imaging and passive acoustic detection systems have fundamentally different availability and detection biases. While passive acoustic monitoring systems depend on the animal to frequently vocalize, thermal imaging systems depend on surfacing events. From a conservation perspective, where we want to maximize our capability to detect animals, both detection approaches can be seen as complementary and should be implemented jointly because they suffer from such different availability biases.

While acoustic detectability of killer whales is dependent on underwater noise levels and thus noise created by dense vessel traffic can mask killer whale calls, TI system detectability is not impacted by underwater noise levels. Furthermore, TI systems do not depend on frequent vocalization to facilitate detection. The benefit of noise independent detection probability of TI systems diminishes during inclement weather conditions, especially with strong wind and rain. This is comparable to shore-based visual marine mammal observation (MMO), where strong wind and high sea states will reduce the visibility of a whale surfacing and a whale blow will be less pronounced and dissipate quickly. In fog and rain conditions visibility in long wave infrared spectrum has been reported to exceed the visibility in the visual spectrum for light to medium fog (factor 5 for Cat1, factor 4 for Cat2) and equal ranges for heavy fog (Cat3) (Beier and Gemperlein, 2004; Flir Teledyne, 2020). Under these conditions, human observers and TI systems do not perform well. A TI system monitors 24/7, does not require someone to be present observing and therefore does not suffer from observer fatigue as MMOs do. A limitation of a TI system is that it cannot distinguish the killer whale ecotype. This is of less concern for vessel-strike mitigation because all whales should be protected from vessel strikes, but it is relevant for management purposes. Passive acoustic monitoring on the other hand can distinguish ecotypes because of differences in acoustic signatures (discrete calls) used by the two ecotypes (Ford, 1989; Deecke, Ford and Slater, 2005). The benefit of integrating several detection methods is the subject of a study that is currently underway and includes the results from this work (Quayle et al. in prep). Ecotype classification during daylight hours could be made possible with the addition of high-resolution visual cameras to the thermal camera systems.

## Conclusion

One of the main goals of this study was to determine if TI is an effective method to detect and localize killer whales including endangered SRKW. The results of this study show that automated TI systems are a viable method for effective long-term monitoring of several marine mammal species including killer whales.

The implied capability to localize each detection provides an opportunity to estimate and continuously update a detection function for different species. It also allows the assessment of collision risks if the location of vessels at the time of the detections are known and to issue alerts to mariners about the possibility, they may encounter whales at specific locations. While species identification capabilities via TI are limited (e.g., thermal detection cannot differentiate between different killer whale ecotypes), the current capabilities are enough to distinguish between species, and ongoing development is likely to improve this capability, at least during the daytime when high-resolution visual data can support the identification. Furthermore, all killer whales in the Salish Sea are considered threatened while SRKW are considered endangered thereby making the TI detection and localization tool an effective conservation tool for killer whales.

Due to the relatively low acquisition cost (< $20k USD) of land-based TI systems, they are a viable option for widespread application at various near-shore locations with high risks of physical disturbance of threatened and endangered whales. The construction of a coastal TI monitoring system network for marine mammal conservation and especially killer whale population recovery is recommended based on the results of this study. A network of TI systems perhaps in combination with other detection methods to detect, track and monitor whale movements in critical high traffic areas can provide near real-time alerts for mitigation measures by competent authorities.

## Acknowledgements

This work is funded by the Whale Detection and Collision Avoidance (WDCA) initiative (Department of Fisheries and Oceans Canada (DFO) G&C WD 18-P-03 and Transport Canada T8009-180317/001/XLV). The system on Saturna Island was possible thanks to the financial support and funding from Marine Environmental Observation Prediction and Response Network (MEOPAR).

We would also like to thank our local partners, BC Ferries (Galliano Island, Active Pass), and the Saturna Island Marine Research and Education Society (SIMRES, Saturna Island, Boundary Pass) for providing access to their infrastructure, power, internet connection and on-site support.

## Author Contributions

Conceptualization: SR, HY, DPZ

Methodology & Data Analysis: SR, AW, DPZ

Field Work: HY, NS, PDH, EC, AW, DPZ

Writing – original draft: SR, HY, DPZ

Writing – review & editing –SR, HY, EC, NS, PDH, AW, DPZ

All authors contributed critically to the drafts and gave final approval for publication.

## Footnotes

1 https://www.canada.ca/en/fisheries-oceans/news/2021/02/government-of-canada-continues-to-invest-in-research-to-inform-protection-measures-for-vulnerable-whale-populations.html

## References

Baille, LMR and Zitterbart, DP (2022) ‘Effectiveness of surface-based detection methods for vessel strike mitigation of North Atlantic right whales’, Endangered Species Research, 49, pp. 57–69.

Baird, R.W. and Dill, L.M. (1996) ‘Ecological and social determinants of group size in transient killer whales’, Behavioral Ecology, 7(4), pp. 408–416. Available at: 10.1093/beheco/7.4.408.

Baird, R.W. and Whitehead, H. (2000) ‘Social organization of mammal-eating killer whales: group stability and dispersal patterns’, Canadian Journal of Zoology, 78(12), pp. 2096–2105. Available at: 10.1139/z00-155.

Beier, K. and Gemperlein, H. (2004) ‘Simulation of infrared detection range at fog conditions for Enhanced Vision Systems in civil aviation’, Aerospace Science and Technology, 8(1), pp. 63–71. Available at: 10.1016/j.ast.2003.09.002.

Buckland, S.T. et al. (2001) ‘Introduction to distance sampling: estimating abundance of biological populations’.

Buckland, S.T. et al. (2004) Advanced Distance Sampling: Estimating abundance of biological populations. OUP Oxford.

Cosewic (2008) ‘COSEWIC assessment and update status report on the killer whale Orcinus orca, Southern Resident population, Northern Resident population, West Coast Transient population, Offshore population and Northwest Atlantic/Eastern Arctic population, in Canada’, Committee on the Status of Endangered Wildlife in Canada, p. viii+–65.

Deecke, V.B., Ford, J.K.B. and Slater, P.J.B. (2005) ‘The vocal behaviour of mammal-eating killer whales: communicating with costly calls’, Animal Behaviour, 69(2), pp. 395–405. Available at: 10.1016/j.anbehav.2004.04.014.

DFO (2018) Southern Resident Killer Whale: imminent threat assessment. Available at: https://www.canada.ca/en/environment-climate-change/services/species-risk-public-registry/related-information/southern-resident-killer-whale-imminent-threat-assessment.html (Accessed: 18 March 2023).

Emmons, C.K., Hanson, M.B. and Lammers, M.O. (2021) ‘Passive acoustic monitoring reveals spatiotemporal segregation of two fish-eating killer whale Orcinus orca populations in proposed critical habitat’, Endangered Species Research, 44, pp. 253–261. Available at: 10.3354/esr01099.

Evans, B.E., Mosby, C.E. and Mortelliti, A. (2019) ‘Assessing arrays of multiple trail cameras to detect North American mammals’, PLOS ONE, 14(6), p. e0217543. Available at: 10.1371/journal.pone.0217543.

Fisheries and Oceans Canada (2011) Recovery Strategy for the Northern and Southern Resident Killer Whales (Orcinus orca) in Canada - Species at Risk Public Registry. Ottawa, p. ix+80 pp. Available at: https://www.sararegistry.gc.ca/document/doc1341a/ind_e.cfm (Accessed: 10 March 2023).

Flir Teledyne (2020) Can Thermal Imaging See Through Fog and Rain? | Teledyne FLIR. Available at: https://www.flir.com/discover/rd-science/can-thermal-imaging-see-through-fog-and-rain/ (Accessed: 10 March 2023).

Ford, J. and Nichol, L. (2014) ‘Marine Mammals of British Columbia’, Victoria, British Columbia: Royal BC Museum. [Preprint].

Ford, J.K. et al. (2017) Habitats of special importance to resident killer whales (Orcinus orca) off the West Coast of Canada. Fisheries and Oceans Canada, Ecosystems and Oceans Science.

Ford, J.K.B. (1989) ‘Acoustic behaviour of resident killer whales (Orcinus orca) off Vancouver Island, British Columbia’, Canadian Journal of Zoology, 67(3), pp. 727–745. Available at: 10.1139/z89-105.

Ford, J.K.B., Ellis, G. and Stredulinsky, E.H. (2013) Information in support of the identification of critical habitat for Transient killer whales (Orcinus orca) off the west coast of Canada. Canadian Science Advisory Secretariat= Secrétariat canadien de consultation ….

Ford, J.K.B. and Ellis, G.M. (2006) ‘Selective foraging by fish-eating killer whales Orcinus orca in British Columbia’, Marine Ecology Progress Series, 316, pp. 185–199. Available at: 10.3354/meps316185.

Graber, J. et al. (2011) ‘Land-based infrared imagery for marine mammal detection’, in Remote Sensing and Modeling of Ecosystems for Sustainability VIII. Remote Sensing and Modeling of Ecosystems for Sustainability VIII, SPIE, pp. 107–117. Available at: 10.1117/12.892787.

Hanson, M. et al. (2010) ‘Species and stock identification of prey consumed by endangered southern resident killer whales in their summer range’, Endangered Species Research, 11, pp. 69–82. Available at: 10.3354/esr00263.

Kalan, A.K. et al. (2016) ‘Passive acoustic monitoring reveals group ranging and territory use: a case study of wild chimpanzees (Pan troglodytes)’, Frontiers in Zoology, 13(1), p. 34. Available at: 10.1186/s12983-016-0167-8.

Lahoz-Monfort, J.J. and Magrath, M.J.L. (2021) ‘A Comprehensive Overview of Technologies for Species and Habitat Monitoring and Conservation’, BioScience, 71(10), pp. 1038–1062. Available at: 10.1093/biosci/biab073.

NOAA (2005) Southern Resident Killer Whale (Orcinus orca) | NOAA Fisheries, NOAA. Available at: https://www.fisheries.noaa.gov/west-coast/endangered-species-conservation/southern-resident-killer-whale-orcinus-orca (Accessed: 18 March 2023).

Palmer, K.J. et al. (2019) ‘Habitat use of a coastal delphinid population investigated using passive acoustic monitoring’, Aquatic Conservation: Marine and Freshwater Ecosystems, 29(S1), pp. 254–270. Available at: 10.1002/aqc.3166.

Parsons, K.M. et al. (2013) ‘Geographic Patterns of Genetic Differentiation among Killer Whales in the Northern North Pacific’, Journal of Heredity, 104(6), pp. 737–754. Available at: 10.1093/jhered/est037.

Rice, A. et al. (2017) ‘Spatial and temporal occurrence of killer whale ecotypes off the outer coast of Washington State, USA’, Marine Ecology Progress Series, 572, pp. 255–268. Available at: 10.3354/meps12158.

Shields, M.W. (2023) ‘2018–2022 Southern Resident killer whale presence in the Salish Sea: continued shifts in habitat usage’, PeerJ, 11, p. e15635. Available at: 10.7717/peerj.15635.

Smith, H.R. et al. (2020) ‘A field comparison of marine mammal detections via visual, acoustic, and infrared (IR) imaging methods offshore Atlantic Canada’, Marine Pollution Bulletin, 154, p. 111026. Available at: 10.1016/j.marpolbul.2020.111026.

Thomas, L. et al. (2010) ‘Distance software: design and analysis of distance sampling surveys for estimating population size’, Journal of Applied Ecology, 47(1), pp. 5–14. Available at: 10.1111/j.1365-2664.2009.01737.x.

Verfuss, U.K. et al. (2016) ‘Low Visibility Real-Time Monitoring Techniques Review’, p. 224.

Verfuss, U.K. et al. (2018) ‘Comparing methods suitable for monitoring marine mammals in low visibility conditions during seismic surveys’, Marine Pollution Bulletin, 126, pp. 1–18. Available at: 10.1016/j.marpolbul.2017.10.034.

Yurk, H. et al. (2010) ‘Sequential Habitat Use by Two Resident Killer Whale (Orcinus orca) Clans in Resurrection Bay, Alaska, as Determined by Remote Acoustic Monitoring’, Aquatic Mammals, 36, pp. 67–78. Available at: 10.1578/AM.36.1.2010.67.

Zitterbart, D.P. et al. (2013) ‘Automatic Round-the-Clock Detection of Whales for Mitigation from Underwater Noise Impacts’, PLOS ONE, 8(8), p. e71217. Available at: 10.1371/journal.pone.0071217.

Zitterbart, D.P. et al. (2020) ‘Scaling the Laws of Thermal Imaging–Based Whale Detection’, Journal of Atmospheric and Oceanic Technology, 37. Available at: 10.1175/JTECH-D-19-0054.1.

