## Supplement for "Coastal Marine Mammal conservation using thermal imaging-based detection systems"

### Supplement Figure 1 | Location

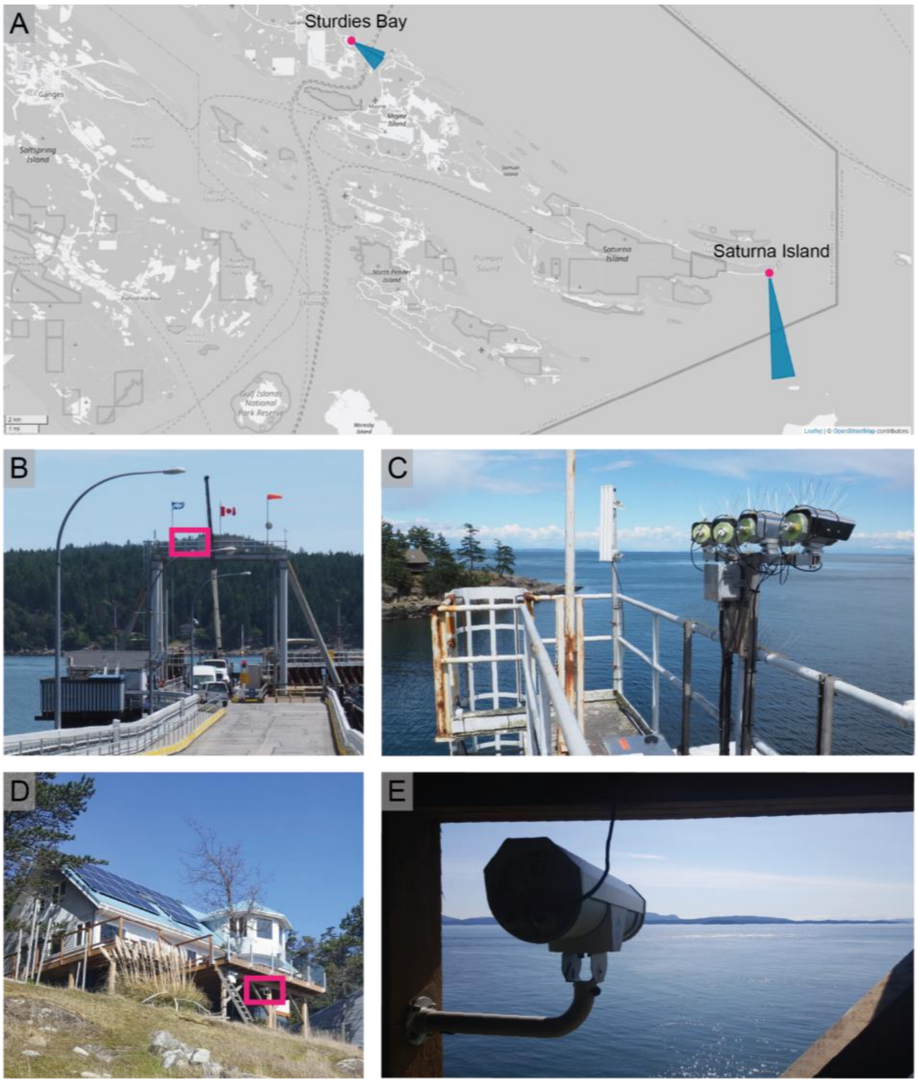


*Supplement Figure **1** | Location of the two deployed systems and their respective field of view (A) Studies Bay (two cameras with 12.5° and 25° horizontal field of view, 14m elevation) and Saturna Island (one camera with 12.5° field of view, 15m elevation). The Sturdies Bay system is installed on top of the Sturdies Bay ferry landing (B) and cameras point across Active Pass (C). The Saturna Island system is installed on a property provided by SIMRES (D) and the camera overlooks Boundary Pass to the South of the island towards Skipjack Island and Waldron Island (D).*
